# Synergistic Effect of Serotonin 1A and Serotonin 1B/D Receptor Agonists in the Treatment of L-DOPA-Induced Dyskinesia in 6-Hydroxydopamine-Lesioned Rats

**DOI:** 10.1101/2021.08.31.458314

**Authors:** Mikael Thomsen, Anca Stoica, Kenneth Vielsted Christensen, Jens Mikkelsen, John Bondo Hansen

## Abstract

**Background:** The gold standard for symptomatic relief of Parkinson’s disease is L-DOPA. However, long-term treatment often leads to motor complications such as L-DOPA-induced dyskinesia (LID). While amantadine (Gocovri™) is the only approved therapy for dyskinesia in people with Parkinson’s disease on the American market, it is associated with neurological side effects and limited efficacy. Thus, there remains a high unmet need for addressing LID in Parkinson’s disease patients worldwide.

**Objective:** The objective of this study was to evaluate the efficacy, safety and performance compared to approved treatments of the serotonin receptor 1A (5-HT1A) and 5-HT1B/D agonists buspirone and zolmitriptan in the 6-hydroxydopamine unilaterally lesioned rat model for Parkinson’s disease.

**Methods:** The hemi-parkinsonian 6-OHDA lesioned rats underwent chronic treatment with L-DOPA to induce dyskinesia and were subsequently used for efficacy testing of buspirone, zolmitriptan and comparison with amantadine, measured as abnormal involuntary movement (AIM) scores after L-DOPA challenge. Safety testing was performed in model and naive animals using forelimb adjusting, rotarod and open field tests.

**Results:** 5-HT1A and 5-HT1B/D agonism effectively reduced AIM scores in a synergistic manner. The drug combination of buspirone and zolmitriptan was safe and did not lead to tolerance development following sub-chronic administration. Head-to-head comparison with amantadine showed superior performance of buspirone and zolmitriptan in the model.

**Conclusions:** The strong anti-dyskinetic effect found with combined 5-HT1A and 5-HT1B/D agonism renders buspirone and zolmitriptan a potential clinical candidate for LID in Parkinson’s disease.

## 1. Introduction

Parkinson’s disease (PD), affecting approximately 1 in 100 people over the age of 60, is a devastating neurological disorder historically characterized by the pathological hallmarks of accumulated protein inclusions known as Lewy bodies and neurites ^1,2^ as well as the progressive loss of the dopamine-producing neurons in the *substantia nigra pars compacta* (SNc). In consequence, the concomitant loss of dopamine leads to profound changes in dopamine-mediated neurotransmission in the basal ganglia network that controls motor coordination in humans^3^. Symptomatic relief of the motor symptoms in Parkinson’s disease patients is currently achieved with levodopa (L-DOPA), a precursor of dopamine, which in the brain is taken up by not only dopamine-producing cells but also serotonergic neurons projecting from the dorsal raphe nucleus, and converted by dopa decarboxylase into the neurotransmitter dopamine that subsequently is released to restore dopamine-mediated neurotransmission^4^.

Under normal conditions the extracellular dopamine concentration in the striatum is finely balanced by presynaptic uptake of dopamine by the dopamine transporter as well as autoregulatory inhibition via the dopamine D2 receptor ^5^. Under the progression of PD and the concomitant loss of dopamine producing cells, the feedback loop is gradually lost whereas serotonergic neurons are still able to take up L-DOPA, convert it to dopamine and release it in a pulsative fashion that leads to an overactivation of the entire basal ganglia network^6^. Consequently, use of L-DOPA in advanced stages of the disease is associated with dysregulated dopamine neurotransmission that gives rise to a hyperkinetic state characterized by abnormal involuntary movements known as L-DOPA induced dyskinesia (LID)^7^. It is considered that LID requires severe loss of nigrostriatal dopamine neurons, a pulsatile delivery of dopamine from serotonergic neurons in the dorsal raphe nuclei and a well-preserved postsynaptic nigrostriatal system in the striatum^8^. L-DOPA-induced dyskinesia is manifest, and more than half of the patients treated with L-DOPA develop dyskinesia within the first five years of therapy ^9,10^. LID frequency increases significantly with higher L-DOPA doses, as well as with young age of PD onset, and disease duration ^11^.

The weak NMDA receptor antagonist amantadine can reduce dyskinesia intensity, but with limited efficacy and associated with adverse effects (such as dizziness, insomnia, hallucinations) at the high doses needed for effective treatment of LID, and tolerance development has also been reported ^12^. An extended-release form of the drug (Gocovri™, Adamas Pharmaceuticals, USA) is currently approved for treatment of dyskinesia in Parkinson’s disease patients, with superior peak plasma concentrations and with sustained levels throughout the day compared to the amantadine immediate release formulation that has been prescribed as off-label treatment for dyskinesia.

Serotonin neurons are equipped with the same enzymatic machinery as dopaminergic neurons to convert L-DOPA to dopamine, store it and release it in the extracellular medium, however they lack the feedback mechanisms as evidenced by Sellnow et al.^13^, to control its synaptic levels and thereby avoid the excessive stimulation of dopamine receptors ^14^. In addition, serotonin neurons can take up dopamine from the extracellular medium via the serotonin transporter (SERT)^15^ and are expected to gradually take over the conversion and the pulsative release of DA, leading to a pathological state with excessive dopamine receptor activation that is likely responsible for L-DOPA-induced dyskinesia. PD patients suffering from LID have a relatively preserved serotonergic system that contributes to the abnormal increases in synaptic dopamine following L-DOPA administration ^16^. Furthermore, LID-affected patients present high serotonergic to dopaminergic terminal availability ratio ^17^ as well as an increase in serotonin transporter to dopamine transporter ratio with PD progression ^18^, thus supporting the involvement of the serotonergic system in the molecular mechanisms of Parkinson’s disease and L-DOPA-induced dyskinesia.

Notably, projection neurons from the dorsal raphe are not regulated by an autoregulatory feedback via dopamine D2 receptors, as nigral dopaminergic cells, but by serotonin 5-HT1A and 5-HT1B receptors^19,20^. Thus, serotonin 5-HT1A and 5-HT1B receptors, either individually or combined, have been considered as targets for LID and studied in PD animal models. Activation of serotonin 5-HT1A receptors using the selective agonist 8-OH-DPAT in animals with a compromised presynaptic nigrostriatal system attenuates the increase in striatal extracellular dopamine derived from administration of exogenous L-DOPA ^21^, suggesting that such agonists could be used to regulate the concentration of dopamine following L-DOPA administration. There is abundant evidence from animal model studies that serotonin receptor modulation with 5-HT1A agonists such as 8-OH-DPAT ^5,22–24^, buspirone ^25,26^, piclozotan ^27^, sarizotan ^28,29^ can reduce LID symptoms. Likewise, evidence from exploratory clinical trials supports anti-dyskinetic effects of the 5-HT1A agonists in Parkinson’s disease patients on L-DOPA treatment^16,30–32^. High 5-HT1A intrinsic agonistic activity is associated with risk of adverse events including a worsening of PD motor complication ^32,33^ thus impacting the therapeutic potential of 5-HT1A agonists. This is hypothesized to be mediated either through activation of pre-synaptic 5-HT1A auto-receptors or by off-target activities on dopamine receptors^34–37^.

Since 5-HT1B receptors have been shown to impact neurotransmitter release in the striatum^38–40^ the therapeutic index limitations from 5-HT1A agonism might be overcome by combining lower and safe doses with a simultaneous effect coming from 5-HT1B agonists acting at the same presynaptic terminals.

Compatible with this concept studies suggest that a combination of 5-HT1A and 5-HT1B agonists could act synergistically and be a better treatment that drugs acting on only one of the receptors. Accordingly, LID studies in PD model monkeys ^41^ and rats ^5^ shows that the combination was effective at doses that were individually ineffective at completely reducing LID. This supports using the modulation of other neurobiological targets important for the development of dyskinesia to improve the efficacy and therapeutic profile of a 5-HT1A agonist.

Another way of impacting the hyperkinetic disease state is to restore the imbalance in the entire basal ganglia network by reducing glutamatergic output from cortical-striatal projection neurons onto the striatal medium spiny neurons. Serotonin 5-HT1D receptors in the human brain affect glutamate release ^42^ and there is data suggesting that the 5-HT1D and the 5-HT1B receptors can be differentiated on the basis of their ability to modulate release of glutamate (5-HT1D) or serotonin (5-HT1B) ^43–45^.

Similarly, balancing the GABAergic output from the striatal medium spiny neurons of the indirect (iMSNs) and direct (dMSN) pathway impacts the hyperkinetic state in LID PD. Preclinical and clinical evidence for such a postsynaptic mechanism is observed with the NMDA-R antagonist amantadine^46^. Amantadine mediates the effect by reducing the postsynaptic activation of NMDA-Rs on both iMSNs and dMSNs^47^; however, since the NMDA-R is broadly expressed in the brain amantadine is also associated with a number of serious adverse events such as constipation, orthostatic hypotension and edema, neuropsychiatric symptoms such as visual and auditory hallucinations, anxiety, nausea and livedo reticularis that may limit the use of amantadine in the PD patient population^48^. Postsynaptic activation of 5-HT1D receptors inhibits adenylate cyclase mediated increases cAMP levels^49,50^. In the MSNs where 5-HT1D receptors are selectively expressed^51^ activation in the dyskinetic disease state could theoretically lead to a decreased neuronal excitability.

We therefore hypothesized that the simultaneous agonism of 5-HT1A and 5-HT1B/D receptors could alleviate LID. To test this hypothesis, we investigated the effect of the 5-HT1A agonist buspirone in combination with the 5-HT1B/D agonist zolmitriptan on L-DOPA-induced dyskinesia in the 6-OHDA-lesioned hemi-parkinsonian rat model.

## 2. Materials and methods

### 2.1. Animals

Experimentally naïve male Sprague-Dawley rats weighing ≈250 g (Shanghai SLAC Co. Ltd.) were housed in groups of 2/cage, with *ad libitum* access to standard rodent chow and water. The animals were housed under controlled environmental conditions, on a 12-hour light-dark cycle. All animal experiments were performed under the Regulations for the Administration of Affairs Concerning Experimental Animals, approved by the State Council of the People’s Republic of China on October 31, 1988.

### 2.2. Unilateral 6-OHDA lesions

The rats were anesthetized with pentobarbital sodium (purchased from the Chinese domestic market) 50 mg/kg (IP), positioned in a stereotaxic frame and injected 6-OHDA (Sigma Aldrich) into the medial forebrain bundle at the following coordinates relative to bregma and dural surface, in mm: tooth bar position -3.3, AP = −1.8, ML = -2, DV = -8.6 (18ug 6-OHDA). Two weeks after surgery the animals were injected apomorphine HCl (Sigma Aldrich, given IP) and contralateral full body turns were recorded over 30 minutes. Only rats with rotational scores ≥180 turns/30 minutes were selected for the next experimental stage.

To generate L-DOPA-induced dyskinesia, one day after the apomorphine-induced rotation test the rats were administered daily 6 mg/kg L-DOPA (Sigma Aldrich) and 15 mg/kg benserazide HCl (purchased from the Chinese domestic market), hereafter denoted as L-DOPA, for 21 days. Thereafter, the rats that had not developed dyskinesia were excluded from the study, and the rats with total axial, limb, orolingual and locomotive abnormal involuntary movement (AIM) scores ≥28 points/120 minutes were kept on a treatment regimen of at least two injections of L-DOPA per week to maintain stable AIM scores. The rats with stable AIM scores were allocated to groups balanced with respect to AIM severity and treated with the drugs or drug combinations as described in the figure legends.

### 2.3. AIM score

AIM score ratings were performed by an investigator not aware of the treatment as described elsewhere ^52^. Briefly, axial (Ax), limb (Li), orolingual (OL) and locomotive AIM were scored on a severity scale from 0 to 4, where 0 = absent, 1 = present during less than half of the observation time, 2 = present for more than half of the observation time, 3 = present all the time but suppressible by external stimuli, and 4 = present all the time and not suppressible by external stimuli.

Although rated in the experiments and included in the model stability evaluations, locomotive AIM scores were excluded from the pharmacology analyses as it was previously established that they do not provide a specific measure of dyskinesia, but instead are an indirect measure of contralateral turning behaviour in 6-OHDA-lesioned rodents (Lundblad et al., 2002, 2005). Raw data were presented as time courses of AIM ratings and integrated AIM scores were calculated from the raw data as area under curve (AUC) using the following formula:

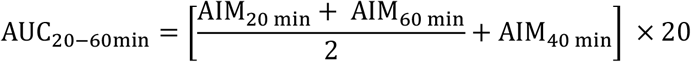

### 2.4. Forelimb adjusting steps

The stepping test ^53–55^ was performed as described previously ^25^ with few modifications. The test is a valid assessment of drug effects on the Parkinson’s disease state, a check of whether the drug combination is affecting the Parkinson’s disease state itself (which could be a side effect in case of worsening or efficacy in case of improvement).

The 6-OHDA-lesioned rats had been kept on at least twice a week chronic treatment with 6 mg/kg L-DOPA combined with 15 mg/kg benserazide HCl to maintain dyskinesia, evaluated for baseline AIM severity and allocated randomly to groups balanced with respect to the baseline test. The baseline adjusting step test consisted of two tests taken on the same day, the mean of which was calculated as baseline.

To evaluate the effect of acute treatment with buspirone HCl (Sigma Aldrich) and zolmitriptan (Damas-Beta) on Parkinson’s disease state, the animals were administered the drugs on the day of testing. For determining the effect of sub-chronic buspirone and zolmitriptan treatment, the animals were administered combinations of L-DOPA, buspirone and zolmitriptan once daily for 3 weeks and the stepping test was performed on the 21^st^ day.

During the stepping test, the rats were held with one forepaw touching the table, and then moved slowly sideways (5 sec for 90 cm) by the experimenter, first in the forehand direction (defined as compensating movement toward the body) and then in the backhand direction (defined as compensating movement away from the body). The number of adjusting steps was counted for both forepaws in the backhand and forehand directions of movement.

Data was presented as percentage of adjusting step of impaired forepaw to intact forepaw and it indicates the degree of forepaw disability.

### 2.5. Rotarod

The rotarod test served the purpose of detecting potentially deleterious effects of the studied compounds on the rats’ motor performance and coordination. In brief, the animals (Sprague Dawley male rats at 9 weeks of age) were trained twice a day for a 3-day period. The rats were placed on the accelerating rod apparatus (Shanghai Jiliang, China) at an initial speed of 4 rotations per minute (rpm), with the speed increasing gradually and automatically to 40 rpm over 300 seconds. Each training trial was ended if the animal fell off or gripped the device and spun around for two consecutive revolutions. The time that rat stayed on the rotarod was recorded. The staying duration recorded at last training trial was used as baseline. Rats were grouped according to a randomly distributed baseline.

For the test session on the fourth day, the rats were evaluated on the Rotarod with the same setting as above 30 min after drug dosing. Dosing and rotarod measurements were conducted by two different scientists. The rotarod performance is expressed as total number of seconds spent on the accelerating rod.

### 2.6. Open field test

The open field test was used to determine the drugs’ effects on locomotor activity. The rats were put in open-field chambers (40 × 40 × 40 cm) 30 minutes after dosing. After a 15-minutes habituation, locomotion was recorded and analysed by Enthovision Video Tracking Software (Noldus Information Technology, Netherlands) for 60 minutes. All locomotor activities were done during dark phase and to eliminate olfactory cues, the arena was thoroughly cleaned with 70% v/v ethanol between tests. The locomotor activity is expressed as total moved distance (cm) every 10 minutes.

### 2.7. Data analysis

Data analysis and statistical tests were carried out using GraphPad Prism 9 software (San Diego, CA, USA). One-way or two-way ANOVA followed by post hoc tests was employed as indicated in the figure legends. Data is presented as mean ± SEM and P < 0.05 was considered statistically significant.

## 3. Results

### 3.1. Buspirone and zolmitriptan reduce LID in 6-OHDA-lesioned rats

The 6-OHDA-leasioned rat is a reliable model for Parkinson’s disease and can be induced to develop dyskinesia upon chronic treatment with L-DOPA. As previously noted, and as shown here, the model is ideally suited for the pharmacological validation of anti-dyskinetic therapies using the AIM score rating scale ^56,57^. For this study, the 6-OHDA-leasioned rat model was validated based on previous studies ^52^, with some modifications (Figure 1A). To evaluate the model validity, after 3 weeks of daily L-DOPA administration, AIM scores were monitored following acute L-DOPA administration (see Figure 1B for a time course of AIM scores during 120 minutes of rating). The results show that the effect of L-DOPA on AIM scores is stable within at least 100 minutes of the observation period (Figure 1B). The animals with ≥28 points/120 minutes were deemed stable PD LID models and continued a twice-weekly L-DOPA administration regimen to maintain dyskinesia.

**Figure 1.**
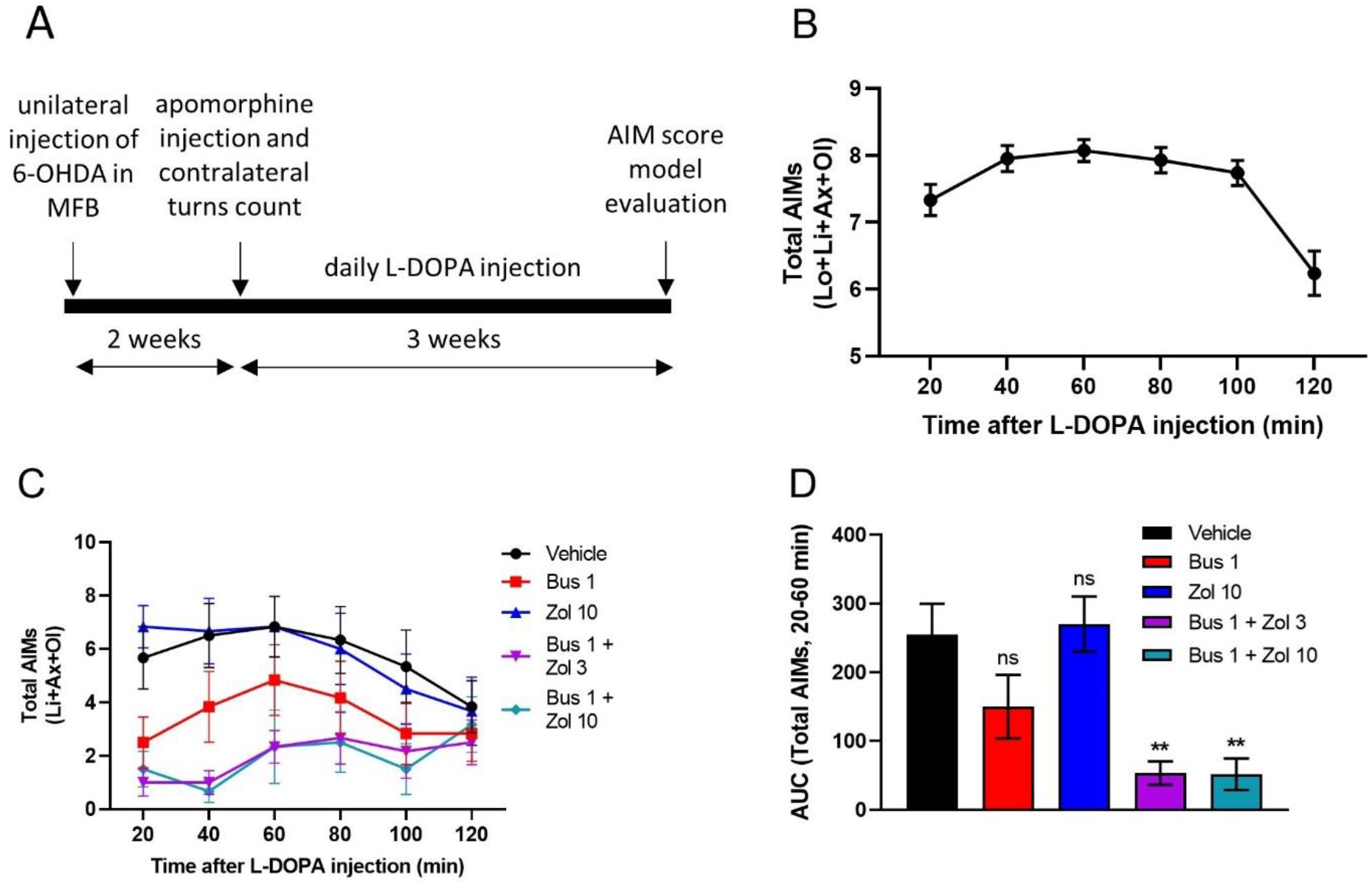
Buspirone and zolmitriptan combined effectively reduce dyskinesia in a PD LID rat model. Schematic representation of the unilateral 6-OHDA-lesioned rat PD model generation (A) and validation (B). Time-course (C) and AUC (D) of total AIM scores following acute IP injection of 1 mg/kg buspirone (1 mg/kg), zolmitriptan (10 mg/kg), buspirone and zolmitriptan (1 mg/kg and 3 or 10 mg/kg, respectively) in 6-OHDA-lesioned PD LID model rats 30 minutes before L-DOPA and benserazide (6 and 15 mg/kg, respectively, injected SC 20 minutes before AIM score rating). Vehicle represents a group treated with L-DOPA and saline. **p<0.01, one-way ANOVA followed by Dunnett’s post-hoc test. Data is presented as mean ± SEM, n=42 (B) and n=6 (C, D).

The effects of buspirone and zolmitriptan alone and in combination on the dyskinesia state in 6-OHDA-lesioned rats were then evaluated after a single administration of L-DOPA (dose SC) by rating the AIM scores. The time course of AIM scores reveals buspirone and zolmitriptan combined suppress the L-DOPA-induced AIM robustly and consistently compared to zolmitriptan alone or the vehicle control (Figure 1C).

The AUC of the 20-60 minutes period after L-DOPA administration shows that 1 mg/kg buspirone in combination with both doses of zolmitriptan (3 or 10 mg/kg) significantly reduced L-DOPA-induced dyskinesia compared to the vehicle control (53 ± 17 AIM and 52 ± 23 AIM respectively, compared to 255 ± 45 AIM, P < 0.01, Figure 1D; statistical significance determined by one-way ANOVA followed by Dunnett’s test, data reported as mean ± SEM, n=6), while buspirone as mono treatment had a nominal but shorter-lived and statistically insignificant effect compared to vehicle treated animals (150 ± 46 AIM compared to 255 ± 45 AIM, mean ± SEM, n=6, p=0.15) and zolmitriptan alone had no effect at all (270 ± 40 compared to255 ± 45 AIM, mean ± SEM, n=6, p=0.99). Collectively, these data show that combining buspirone and zolmitriptan leads to synergistic effect in reducing L-DOPA-induced dyskinesia in 6-OHDA-lesioned rats.

### 3.2. Sub-chronic treatment with buspirone and zolmitriptan does not cause tolerance development

To determine whether long-term combined treatment with buspirone and zolmitriptan leads to tolerance development, dyskinetic 6-OHDA-lesioned rats were kept on a 3-week daily treatment regimen of either L-DOPA alone, referred to as scL, or L-DOPA co-administered with buspirone (0.5 mg/kg) and zolmitriptan (10 mg/kg), referred to scLBZ, as described in Figure 2A. After the sub-chronic treatment, the animals were split further into 4 groups and were acutely administered L-DOPA alone or L-DOPA co-administered with buspirone (0.5 mg/kg) and zolmitriptan (10 mg/kg), after which the animals were evaluated for AIM scores (Figure 2A).

**Figure 2.**
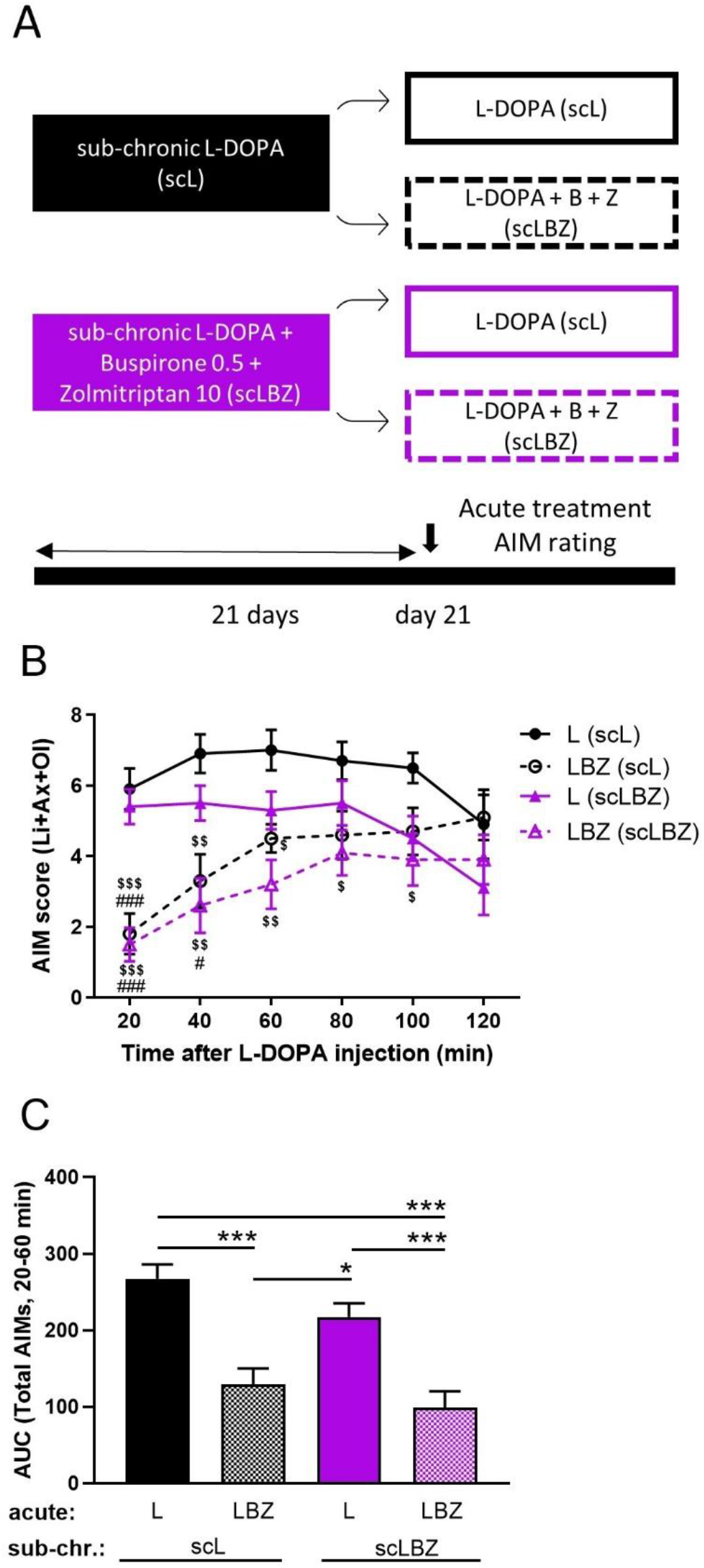
Sub-chronic treatment with buspirone and zolmitriptan does not lead to tolerance development. Schematic representation of the treatment regimen for sub-chronic administration of L-DOPA and benserazide (6mg/kg and 15 mg/kg, respectively, SC, injected 20 minutes before AIM rating) ± buspirone and zolmitriptan (0.5 mg/kg and 10 mg/kg, respectively, IP, 30 minutes before AIM rating) in 6-OHDA-lesioned PD LID rats (A). Time-course (B) and AUC (C) of total AIM scores following acute administration of the same doses of L-DOPA and benserazide ± buspirone and zolmitriptan in the sub-chronically treated rats. *p<0.05, ***p<0.001, one-way ANOVA followed by Tukey’s post-hoc test. ^$^p<0.05, ^$$^p<0.01; ^$$$^p<0.001 vs. L(scL); ^#^p<0.05, ^###^p<0.001 vs. L(scLBZ), two-way ANOVA followed by Tukey’s post-hoc test. Data is presented as mean ± SEM, n=10.

The results show a 0.5-fold change in AIM scores following acute treatment with buspirone and zolmitriptan both in rats pre-treated for 3 weeks with buspirone and zolmitriptan and in animals pre-treated for 3 weeks with L-DOPA alone (Figure 2B and C, compare LBZ (scLBZ) with LBZ (scL)), indicating that sub-chronic treatment with a combination of buspirone and zolmitriptan does not cause the development of tolerance. There was no significant difference between the acute L-DOPA-treated groups (Figure 2B and C, L(scL) compared to L(scLBZ)), indicating that a 3-week-long treatment with buspirone and zolmitriptan did not prevent L-DOPA induced dyskinesia.

This demonstrates that a combined sub-chronic treatment with buspirone and zolmitriptan in L-DOPA-induced dyskinesia model rats is not causing the development of significant tolerance with the risk of a deterioration of the treatment effect.

### 3.3. Acute or sub-chronic treatment with buspirone and zolmitriptan does not affect Parkinson’s disease symptoms or the coordination and locomotor activity of rats

Serotonergic treatments that attenuate dyskinesia can interfere with the anti-akinetic effects of L-DOPA in both animal models and humans ^33,58–60^. To assess whether the combination of buspirone and zolmitriptan is affecting classical motor symptoms of PD or the efficacy of L-DOPA treatment, 6-OHDA-lesioned hemi-parkinsonian rats were subjected to the forelimb adjusting step test following acute and sub-chronic buspirone and zolmitriptan combination treatment. The forelimb adjusting step test serves to detect motor initiation deficits in the forelimbs of lesioned rats ^55^.

The results show that acute treatment with buspirone in combination with zolmitriptan has comparable effects to L-DOPA alone and both perform significantly better than the vehicle-treated group in this model (Figure 3A p<0.01 and 0.05 respectively, one-way ANOVA followed by Dunnett’s test). Following 3 weeks of daily treatment, L-DOPA alone and L-DOPA plus buspirone in combination with zolmitriptan have similar beneficial effects on the percentage of adjusting steps, and both perform better than the untreated (vehicle) group in this model (Figure 3B). The differences between the treatment with vehicle and L-DOPA alone or vehicle and L-DOPA plus the buspirone and zolmitriptan combination are statistically significant (Figure 3B, p<0.001, one-way ANOVA followed by Tukey’s test) whereas there is no statistically significant difference between L-DOPA alone and L-DOPA plus the combination treatment (Figure 3B, p=0.92). A nominal effect of the combination treatment alone is observed but the difference is not statistically significant from the vehicle group (Figure 3B, p=0.26).

**Figure 3.**
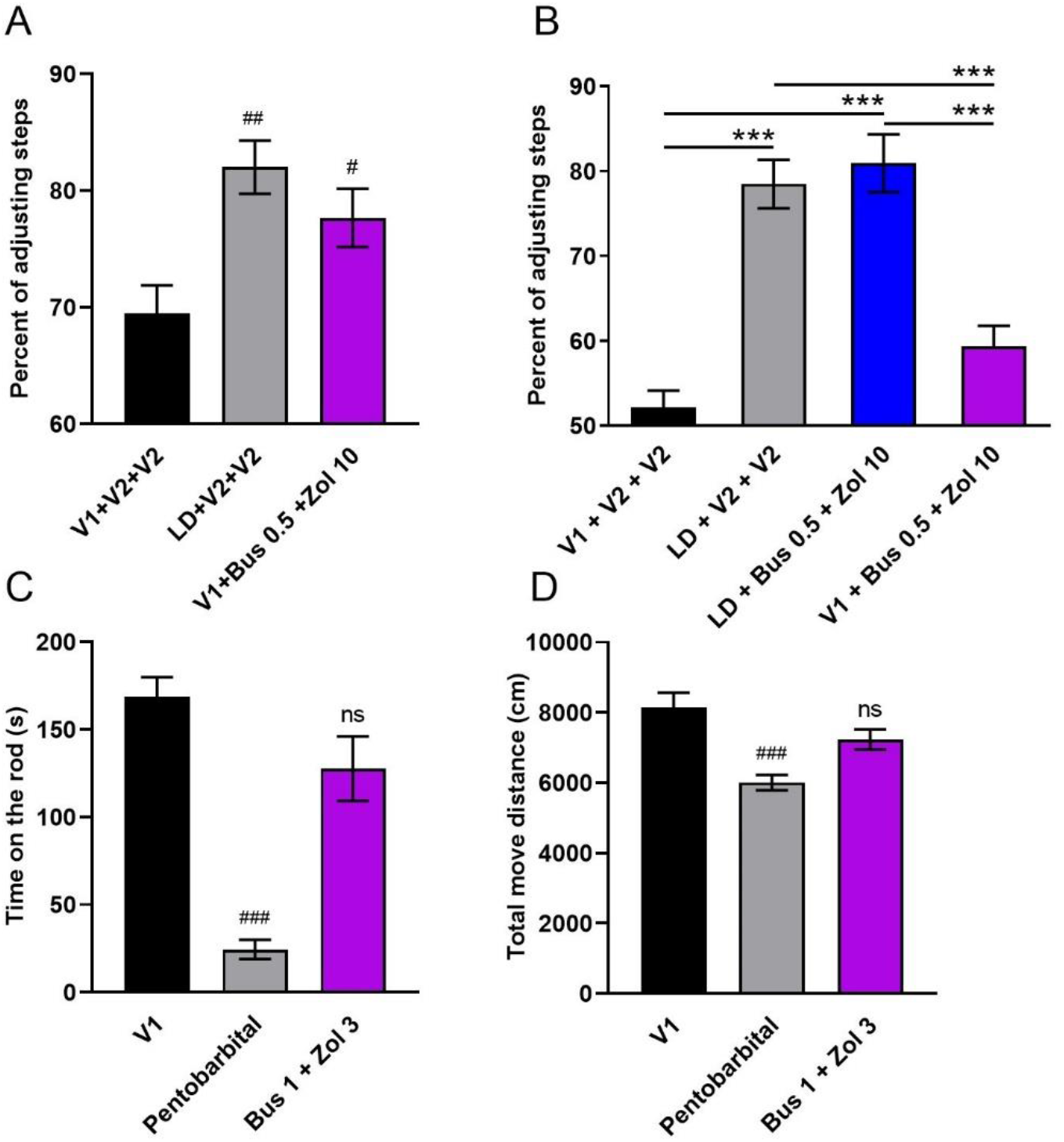
Safety evaluation of the treatment with buspirone and zolmitriptan. The percentage of adjusting steps of impaired forepaw to intact forepaw following acute treatment with L-DOPA and benserazide (3 mg/kg and 15 mg/kg, respectively, SC, 60 minutes pre-test) or with buspirone and zolmitriptan (0.5 mg/kg and 10 mg/kg, respectively, IP, 30 minutes pre-test) in 6-OHDA-leasioned PD LID rats showed that buspirone and zolmitriptan do not have negative effects on Parkinson’s disease symptoms (A). Daily 3-week treatment with L-DOPA and benserazide (3 mg/kg and 15 mg/kg, respectively, SC, 60 minutes pre-test) alone, in combination with buspirone and zolmitriptan (0.5 mg/kg and 10 mg/kg, respectively, IP, 30 minutes pre-test) or with buspirone and zolmitriptan alone in in 6-OHDA-leasioned PD LID rats showed that buspirone and zolmitriptan do not interfere with the anti-akinetic effects of L-DOPA (B). The time spent on the rotarod (C) and total move distance in the open field test (D) of naïve rats treated with buspirone and zolmitriptan (1 mg/kg and 3 mg/kg, respectively, IP, 30-35 minutes pre-test) compared to rats treated with pentobarbital (15 mg/kg, IP, 30 minutes pre-test) revealed that buspirone and zolmitriptan combined do not have sedative effects. V1 represents saline, V2 represents 10% Tween-80. ***p<0.001, one-way ANOVA followed by Tukey’s post-hoc test. ^#^p<0.05, ^##^p<0.01, ^###^p<0.001 vs. Vehicle control, one-way ANOVA followed by Dunnett’s post-hoc test. Data is presented as mean ± SEM, n=9-12.

This demonstrates that the combination of buspirone and zolmitriptan, in the current setting and at doses effective in treating L-DOPA induced dyskinesia in the rat, is unlikely to have any negative effects on the Parkinson’s disease state and the anti-akinetic effect of L-DOPA.

The rotarod test was used to detect treatment-induced motor deficits in normal rats, and the sedative barbiturate pentobarbital served as positive control. Animals treated with buspirone in combination with zolmitriptan had no statistically significant different performance in the rotarod model compared to vehicle treated animals, showing that motor performance and coordination are not significantly impaired in rats after administration of the combination (Figure 3C, p=0.06, one-way ANOVA followed by Dunnett’s test). As expected, pentobarbital reduced the time rats spent on the rotarod when compared to the vehicle treated group; the difference was statistically significant (Figure 3C, p<0.001).

The open field test was used to determine the effects of buspirone and zolmitriptan on locomotor activity in rodents. The results revealed that compared to the vehicle buspirone in combination with zolmitriptan had no statistically significant effects on rat performance in the open field test measured as total moved distance during a 30-minute observation period (Figure 3D, p=0.1, one-way ANOVA followed by Dunnett’s test). Pentobarbital significantly reduced rat motor performance during the total observation period when compared to vehicle treated animals (Figure 3D, p<0.001).

### 3.4. The combined effect of buspirone and zolmitriptan is superior to amantadine in reducing AIMs

We compared the effect of the NMDA-R antagonist amantadine at doses selected based on published data ^52^ as well as a dose-range-finding study (data not shown), with the effect of buspirone and zolmitriptan in combination in 6-OHDA-lesioned LID model rats.

Treatment with buspirone (0.5 mg/kg or 1 mg/kg) and zolmitriptan (10 mg/kg) had statistically significant inhibitory effects on L-DOPA induced dyskinesia in terms of AUC of total AIM scores when compared to the vehicle group (Figure 4). While amantadine (40 or 60 mg/kg) showed minor nominal effects in reduction of AIM scores, but not statistically significant (Figure 4B, p=0.17 and p=0.09 for 40 and 60 mg/kg amantadine, respectively; one-way ANOVA followed by Tukey’s test). Both doses of buspirone (0.5 mg/kg, or 1mg/kg) combined with zolmitriptan had statistically significant effects compared to both doses of amantadine tested. The inhibitory effect of buspirone (0.5 mg/kg or 1 mg/kg) combined with zolmitriptan (10 mg/kg) on LID showed a clear dose-dependency.

**Figure 4.**
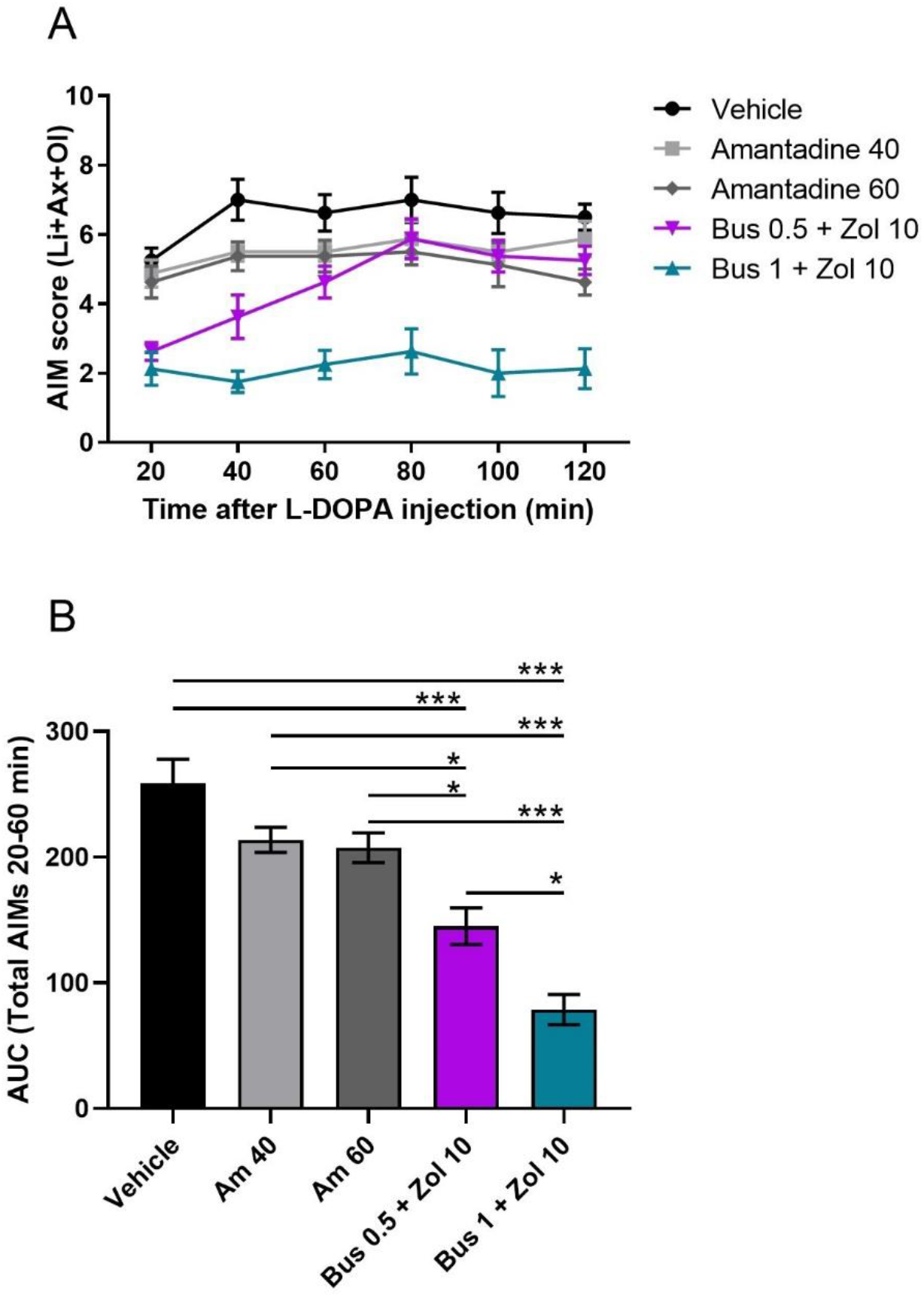
Buspirone and zolmitriptan are significantly more efficacious in reducing AIM scores than amantadine. Time-course (A) and AUC (B) following administration of amantadine (40 or 60 mg/kg, SC, 2h pre-test) and buspirone combined with zolmitriptan (0.5 or 1 mg/kg and 10 mg/kg, respectively, IP, 30 min pre-test) show that both dose combinations of buspirone and zolmitriptan are significantly better at reducing AIM scores than amantadine and the effect is dose-dependent. Vehicle represents 10% Tween-80. ***p<0.001, *p<0.05, one-way ANOVA followed by Tukey’s post-hoc test. Data is presented as mean ± SEM, n=8.

Collectively, in this rat PD dyskinesia model which is predictive for the clinical efficacy of amantadine and under the current test conditions, the combination of buspirone and zolmitriptan is superior to the approved treatment for LID, amantadine.

## 4. Discussion

It has been previously shown that combined 5-HT1A and 5-HT1B agonism potently and safely alleviates LID in 6-OHDA-lesioned rats and MPTP-treated macaques^5,41^, but in the present study we report for the first time that a combination of 5-HT1A and 5-HT1B/D agonism presents with a strong anti-dyskinetic effect in a rodent model of PD dyskinesia. We also show that the anti-dyskinetic effect obtained with the combination of buspirone and zolmitriptan is synergistic of nature: treatment with buspirone or zolmitriptan alone resulted in either a small, statistically insignificant effect or no effect at all, respectively. However, the combination treatment reduced, with up to 80%, the AUC of total AIM scores compared to the vehicle control group.

Moreover, we demonstrate that the combination of buspirone and zolmitriptan does not induce motor complications in L-DOPA-treated hemi-parkinsonian rats. Higher doses of buspirone (2-4 mg/kg) than those used in the present study, while effective at reducing AIM scores, have been shown to cause significant reductions in the rotarod performance of rats ^52^ or to trigger serotonin syndrome symptoms^61^. Eltoprazine, another 5-HT1A agonist, is highly effective in suppressing dyskinesia, but associated with a worsening of the therapeutic effect of L-DOPA^58,62^. The lack of motor complications observed with the combination of buspirone and zolmitriptan may be due to the lower buspirone doses used, which supply sufficient 5-HT1A receptor occupancy to give a strong anti-dyskinetic effect when combined with zolmitriptan, but low enough to circumvent motor side effects. The anti-dyskinetic effect of buspirone was shown to correlate with normalized striatal D1 receptor signalling via normalization of DARPP32 and ERK2 phosphorylation ratios^63^. It is reasonable to speculate that a reduction in cAMP in the dMSN by zolmitriptan could add to the beneficial effect of buspirone on LID. Furthermore, modulation of glutamate release via the 5-HT1D receptor^42^ could contribute to the efficacy of the combination treatment. Tolerance development is a concern when developing new therapies, and can contribute to the high failure rate in translating preclinical to clinical efficacy in neuroscience^64^. We have shown here that the combination of buspirone and zolmitriptan does not lead to tolerance confirming the lack of clinical evidence on development of tolerance for either buspirone or zolmitriptan.

Lastly, we show that in terms of preclinical efficacy the combination is superior to amantadine, the only medication approved for treatment of dyskinesia in Parkinson’s disease patients on L-DOPA therapy. We have tested buspirone and zolmitriptan alongside amantadine and found a ≈44-70% reduction in AUC of total AIM scores relative to the vehicle group, compared to the ≈17-20% reduction observed with amantadine. The doses of amantadine used in the preclinical model (40 and 60 mg/kg) are in line with previous studies^52,65^. Furthermore, the results of a PK/PD correlation analysis of amantadine for LID across multiple species supports that the doses used in this study correlate with the clinically relevant dosage and would ensure an amantadine plasma concentration in rats sufficient for a consistent anti-dyskinetic effect^46^.

To identify novel treatments for motor fluctuations due to LID in Parkinson’s disease an in-depth understanding of the underlying disease biology circuitry in the basal ganglia network is crucial. A pathophysiological model for LID in Parkinson’s disease has been proposed^8^ which postulates the following requirements for LID to manifest in late-stage PD patients: (i) A presynaptic nigrostriatal loss of dopamine neurons in the SNc; (ii) A pulsatile delivery (short half-life) of dopamine from serotonergic neurons projecting from the dorsal raphe nuclei (DRN); and (iii) A relatively well-preserved postsynaptic nigrostriatal system in the striatum, i.e. medium spiny neurons (MSN) of the direct and indirect pathways. Besides this there are other neuronal pathways that can modulate the manifestation of LID such as the cortico-striatal projection neurons that release glutamate in the striatum or the cholinergic neurons that release acetylcholine. In line with this model, the serotonin receptors 5-HT1A and 5-HT1B are highly expressed in the dorsal raphe nucleus^66,67^ and on the cortico-striatal projection neurons^51^, while 5-HT1D expression is enriched in MSN of both the indirect and direct pathway^51^. A decrease in neuronal excitability could theoretically be achieved selectively in the dMSN by 5-HT1D agonism countering the cAMP increase mediated by the G_s/olf_-coupled dopamine D1 receptor^68^, but not in iMSNs where the cAMP levels are already decreased under the inhibition of adenylyl cyclase via the G_i/o_ proteins coupled to the dopamine D2 receptor^69^. Serotonin 5-HT1D receptor agonism could thus be effective in balancing the output from the direct and indirect pathways in a similar way to amantadine, albeit without the safety-related concerns.

In addition it has been shown that 5-HT1B receptor striatal expression is induced in hemi-parkinsonian rats after L-DOPA administration^70^.

This study aimed to identify pre-existing medicines for drug targets localized in the abovementioned neuronal circuitry that would selectively reduce the pulsative dopamine release as well as modulate the imbalance in the entire basal ganglia network. The aim was to achieve this by combining drugs mechanistically acting on presynaptic DRN neurons and/or on pre-synaptic cortico-striatal projection neurons or on postsynaptic MSN neurons. Buspirone, a 5-HT1A receptor partial agonist, and zolmitriptan, a 5-HT1B/D receptors agonist, which are both FDA-approved medicines for anxiety and migraine, respectively, fulfilled those requirements. Activation of 5-HT1A auto-receptors reduces release of serotonin^71,72^ thus it is likely to lead to a reduction in “false neurotransmitter”, i.e. dopamine or glutamate, release from serotonergic neurons and cortical neurons ^16,73^. Similar effects have been shown for 5-HT1B^74,75^. In addition to the effects of 5-HT1A and 5-HT1B, activation of the 5-HT1D receptor, a Gi-protein-coupled receptor, in striatal medium spiny neurons (MSNs) reduces cAMP levels via inhibition of adenylate cyclase^76^ and will lead to decreased cAMP levels in MSNs of both dopamine D1 receptor, direct and dopamine D2 receptor, indirect pathway^69^ thus balancing the basal ganglia output to the thalamus and motor cortex^77^. Collectively, this is hypothesized to underlie the synergistic anti-dyskinetic effect of the buspirone and zolmitriptan combination.

The overall outcome of our studies with the combination of buspirone and zolmitriptan is positive and fulfills the preclinical therapeutic conditions on multiple parameters, including synergy between the two compounds, lack of tolerance development and interference with PD symptoms upon long-term treatment, safety, and performance in comparison to amantadine. Moreover, in the adjusting step test following sub-chronic drug administration, the combination treatment itself without L-DOPA showed a nominal increase in the percentage of adjusting steps compared to the vehicle control group, suggesting buspirone and zolmitriptan might have a beneficial effect on the Parkinson’s disease state. The open field and rotarod test results support that buspirone and zolmitriptan do not have sedative effects and are safe when used at doses that are effective at reducing L-DOPA-induced dyskinesia in rats, in consensus with published reports on safety^78–81^.

There is a high unmet need for addressing motor complications such as wearing off and motor fluctuations to improve the daily life of Parkinson’s disease patients. Around 40-45% of PD patients develop dyskinesia within 5 years of L-DOPA treatment initiation^10,82^, and the number increases to 50-75% after 10 years of L-DOPA treatment^83,84^. PD patients with LID show increased postural instability^85^ and have a higher risk of falls^86^ compared to non-dyskinetic patients. Furthermore LID has a negative social impact and diminishes overall the quality of life of PD patients while increasing the burden on caregivers^87,88^.

Thus, the serotonin receptor agonist combination presented in this study will be developed into a therapy for L-DOPA-induced dyskinesia. Given its synergistic effect primarily on 5-HT1A and 5-HT1B/D, it may show improved efficacy compared to targeting 5-HT1A alone, a strategy that has failed in the past^89^, and to existing medicines. Future clinical studies are necessary to establish the efficacy and safety of this drug combination in PD patients suffering from LID.

